# Cross-tolerance evolution is driven by selection on heat tolerance in *Drosophila subobscura*

**DOI:** 10.1101/2023.09.05.556367

**Authors:** Luis E. Castañeda

**Affiliations:** Programa de Genética Humana, Instituto de Ciencias Biomédicas (ICBM), Facultad de Medicina, Universidad de Chile, Santiago, Chile

**Keywords:** correlated evolution, global warming, heat stress intensity, stress resistance evolution, thermal tolerance landscape

## Abstract

The evolution of heat tolerance is a crucial mechanism for the adaptive response to global warming, but it depends on the genetic variance carried by populations and on the intensity of thermal stress in nature. Experimental selection studies have greatly benefited research into heat tolerance, providing valuable insights into its evolutionary process. However, the impact of varying levels of heat stress intensity on the associated changes in resistance traits has not yet been explored. Here, the correlated evolution of increasing knockdown temperature in *Drosophila subobscura* was evaluated on the knockdown time at different stress temperatures (35, 36, 37, and 38 °C), thermal death time (TDT) curves, and desiccation and starvation resistance. The selection of heat tolerance was performed using different ramping temperatures to compare the impact of heat intensity selection on resistance traits. Correlated evolution was found for these four resistance traits in *D. subobscura*, indicating that the evolutionary response to tolerance of higher temperatures also confers the ability to tolerate other stresses such as desiccation and starvation. However, these correlated responses depended on the intensity of thermal selection and sex, which may limit our ability to generalize these results to natural scenarios. Nevertheless, this study confirms the value of the experimental evolutionary approach for exploring and understanding the adaptive responses of natural populations to global warming.

## INTRODUCTION

Rising environmental temperatures are a major challenge for ectotherms (i.e., organisms whose body temperature depends on the ambient temperature) because their morphology, physiology, behavior, and performance depend on the thermal environment (Huey and Stevenson 1979; Cossins and Bowler 1987; Angilletta 2009). Furthermore, rising environmental temperatures increase the risk of extinction for many species living near their upper thermal limits (Deutsch et al. 2008; Huey et al. 2009; Hoffmann and Sgrò 2011). However, ectotherms can avoid the negative effects of heat through behavioral thermoregulation, evolutionary change, and/or phenotypic plasticity of the upper thermal limits (Visser 2008).

Evolutionary adaptation depends on the genetic variation exhibited by upper thermal limits; however, some studies have suggested that heat tolerance has a limited evolutionary potential to respond to increasing environmental temperatures (Chown et al. 2009; Mitchell and Hoffmann 2010; Kellermann et al. 2012). Yet, theoretical and empirical evidence suggests that heritability estimates for heat tolerance tend to be lower when heat tolerance is measured in longer assays (e.g., slow-ramping assays or static assays using sublethal temperatures) than in shorter assays (e.g., fast-ramping assays or static assays using extremely high temperatures) (Chown et al. 2009; Mitchell and Hoffmann 2010; Rezende et al. 2011; Blackburn et al. 2014; Heerwaarden et al. 2016; Castañeda et al. 2019). Thus, the intensity of heat stress may influence our predictions regarding the evolutionary potential of heat tolerance, but how do populations respond to variable selection driven by heat stress? Selection under laboratory conditions has a long history of providing information on the adaptive evolution of specific selective agents (Lenski and Bennett 1993; Garland Jr 2003; Fuller et al. 2005; Gibbs and Gefen 2009). In particular, the experimental evolution of heat tolerance has been assessed in several species, including fish, corals, and insects (Baer and Travis 2000; Kelly et al. 2012; Geerts et al. 2015; Esperk et al. 2016). Experimental evolution of heat tolerance has also been studied in several *Drosophila* species, *including D. melanogaster* (Gilchrist and Huey 1999; Folk et al. 2006), *D. subobscura* (Quintana and Prevosti 1990; Mesas et al. 2021; Mesas and Castañeda 2023), and *D. buzzatti* (Krebs and Loeschcke 1996). Most of these studies reported the evolution of heat tolerance using fast ramping protocols, ranging from 0.4 °C/min in Folk et al. (Folk et al. 2006) to 1 °C/min in Gilchrist and Huey (Gilchrist and Huey 1999) or static high-temperature assays (40 °C), as in Bubliy and Loeschcke (2005). Recently, Mesas et al. (Mesas et al. 2021) reported that selected lines of *D. subobscura* evolved higher heat tolerance, regardless of the heating rate used during the selection experiments (slow-ramping rate: 0.08 °C/min and fast-ramping rate: 0.4 °C/min).

Interestingly, several of these selection experiments on heat tolerance in *Drosophila* have found correlated responses in other traits such as starvation resistance, desiccation resistance, and heat shock proteins (Hoffmann et al. 1997; Feder et al. 2002; Bubliy and Loeschcke 2005). However, the intensity of thermal stress is expected to have important effects on the correlated responses of other traits to heat tolerance selection (Fragata and Simões 2022). For example, fast-ramping selected lines have evolved thermal performance curves with higher optimum temperatures and narrower thermal breadths than slow-ramping selected lines (Mesas et al. 2021). In addition, Mesas and Castañeda (Mesas and Castañeda 2023) reported that the evolution of heat tolerance was associated with reduced activity of the enzymes involved in the glucose-6-phosphate branch point and increased performance of life-history traits in slow-ramping selected lines. However, they did not observe any changes in the metabolic rate of the selected lines, as predicted by Santos et al. (2012). In summary, there is evidence that heat stress intensity determines the magnitude of the evolutionary responses of performance, metabolic, and life-history traits to heat tolerance selection; however, the correlated evolution of resistance traits has not yet been tested. This information should explain how thermal stress intensity might determine the cross-tolerance evolution to stressful environmental conditions. Natural populations are regularly subjected to multiple environmental stressors, and it is well-established that enhanced tolerance to one stressor can enhance tolerance to another (Rodgers and Gomez Isaza 2023). Cross-tolerance induced by thermal stress has been widely studied in several arthropod species, increasing resistance to desiccation, insecticides, and pathogens (Kalra et al. 2017; Rodgers and Gomez Isaza 2021; Singh et al. 2022). However, the cross-tolerance patterns at the evolutionary level can be constrained or facilitated by genetic correlations among resistance traits depending on the environmental context (Lande and Arnold 1983; Bubliy and Loeschcke 2005; Gerken et al. 2016).

Previous research has examined the impact of varying levels of heat stress on the heat knockdown temperature of *D. subobscura*, as well as its associated impacts on thermal performance curves (Mesas et al. 2021), energy metabolism, and fitness-related traits (Mesas and Castañeda 2023). The evolutionary response of these traits was evaluated using two thermal selection protocols that differed in the rate of temperature increase (hereafter, ramping rate) to measure the heat knockdown temperature: slow-ramping selection (0.08°C min^-1^) and fast-ramping selection (0.4°C min^-1^). The present study investigates the effects of heat intensity selection for increasing knockdown temperature on the cross-tolerance evolution of four different resistance traits in *D. subobscura*: knockdown time at different stress temperatures, thermal-death-time curves (TDT), desiccation resistance, and starvation resistance. In particular, TDT curves represent an integrative approach to assess how the probability of survival depends on the intensity and duration of heat stress, as they allow the estimation of the critical thermal maxima (CT_max_) and thermal sensitivity using the thermal tolerance measurements obtained at different stress temperatures (Rezende et al. 2014). Here, it is expected that fast-ramping selected lines will exhibit higher knockdown time at highly stressful temperatures and higher CT_max_ because fast-ramping protocols reduce the confounding effects (e.g., hardening, rate of resource use) on heat tolerance associated with the assay length (see Rezende et al. 2011; Santos et al. 2012; Mesas et al. 2021). In contrast, slow-ramping selected lines should exhibit higher desiccation and starvation resistance because individuals with higher starvation and desiccation resistance exhibit higher thermal tolerance during long assays.

## Materials and Methods

### Sampling and maintenance

*D. subobscura* females were collected in the spring 2014 at the Botanical Garden of the Universidad Austral de Chile (Valdivia, Chile; 39° 48’ S, 73° 14’ W) using plastic traps containing banana/yeast baits. Two hundred females were collected and placed individually in plastic vials containing David’s killed-yeast *Drosophila* medium to establish isofemale lines. In the next generation, 100 isofemale lines were randomly selected, and 10 females and 10 males per line were placed in an acrylic cage to establish a large, outbred population. In the next generation, the flies from this cage were divided into three population cages (R1, R2, and R3), attempting to assign the same number of flies to each cage. After three generations, the flies in each replicate cage were divided into four population cages, trying to assign the same number of flies to each cage. This procedure established 12 population cages assigned to each artificial selection protocol in triplicate: fast-ramping selection, fast-ramping control, slow-ramping selection, and slow-ramping control lines (Fig. S1). During the selection experiments, population cages were maintained at 18 °C (12:12 light-dark cycle) in a discrete generation, controlled larval density regime (Castañeda et al. 2015). Each population cage had a population size of 1000-1500 breeding adults.

### Heat tolerance selection

For each replicate line, 120 four-day-old virgin females were randomly mated with two males for two days, after which the females were individually placed in a capped 5-mL glass vial, and the males were discarded. The vials were attached to a plastic rack and immersed in a water tank with an initial temperature of 28 °C, controlled by a heating unit (model ED, Julabo Labortechnik, Seelbach, Germany). After an equilibration period of 10 min, the temperature was increased to 0.08 °C min^-1^ for the slow-ramping selection protocol or 0.4 °C min^-1^ for the fast-ramping selection protocol. Assays were stopped when all flies collapsed. Each assay was recorded using a high-resolution camera (model D5100, Nikon, Tokyo, Japan) and then visualized to score the knockdown temperature for each fly, defined as the temperature at which each fly ceased to move. Flies were ranked by knockdown temperature, and four virgin females were selected from the progeny of the 40 flies with the highest knockdown temperature (top 30% of each assay) to establish the next generation. For the fast and slow control lines, the knockdown temperature was measured as described above, but the progeny was randomly selected to establish the next generation, regardless of the knockdown temperature of their mother.

This artificial selection experiment was performed for 16 generations, after which flies from each selection treatment were placed in separate acrylic cages and maintained without selection (e.g., relaxed selection) at 18 °C and a 12:12 light-dark cycle.

### Knockdown time in static assays

Eggs were collected from each population cage and transferred to vials at a density of 40 eggs/vial. At 4 days of age, ten females and ten males from each population cage were tested to measure their heat knockdown time at four different static temperatures: 35, 36, 37, and 38°C. This experimental design allowed the measurement of 960 flies (10 flies × 2 sexes × 4 static temperatures × 4 selection treatments × 3 replicated lines). Static assays were performed similarly to knockdown temperature assays, but static temperatures were used instead of ramping temperatures. A total of 240 flies were measured for each static temperature, except for the assay at 35°C (178 flies) because two flies died before the start of the assay, and a video file of one assay was corrupted (data for 60 flies were lost). For the 37°C assay, four flies died before the assay began, and the collapse time could not be measured for six flies. Finally, for the 38°C assay, three flies died before the start of the assay and the collapse time could not be measured for five flies. Heat knockdown assays were performed in generation 23 (Fig. S1).

### Desiccation and starvation resistance

Eggs from each replicate cage were collected and maintained in vials at a density of 40 eggs/vial. Only fast control lines were measured as control lines. This decision was based on logistical reasons (i.e., the high number of vials) and statistical support because fast and slow control lines did not differ in their knockdown times and CT_max_ values (see *the Results* section).

For desiccation resistance assays, five flies from each sex were separately placed in a vial containing five desiccant droplets (Drierite) and sealed with parafilm (flies had no access to food or water during the assay). For starvation resistance assays, five flies from each sex were separately placed in a vial containing agar only (flies had access to water but no food). For both desiccation and starvation resistance assays, the number of live flies was counted every 3 h until all the flies were dead. Desiccation and starvation resistance were measured in 126 vials containing 10 flies each, respectively (7 vials × 2 sexes × 3 selection treatments × 3 replicate lines). These experiments were conducted at 18 °C using flies from generation 24 (Fig. S1).

### Statistical analysis

Normality and homoscedasticity were tested for all variables, and the knockdown times were squared root transformed to meet the parametric assumptions. All analyses were performed with R software (R Development Core Team 2011).

### Heat tolerance

For the knockdown temperature, control and selection lines were compared separately for the fast- and slow-ramping selection because it is well known that the knockdown temperature is higher in fast-ramping than in slow-ramping assays (Chown et al. 2009; see Mesas et al. 2021). For the knockdown time analysis, a mixed linear model with ramping selection (fixed effect with fast-control, slow-control, fast-selection, and slow-selection lines as levels), sex (fixed effect with females and males as levels), and replicate lines nested within the thermal selection (random effect with replicates 1, 2 and 3 as levels) was performed using the library *lme4* package for R (Bates et al. 2015). Fixed effects were tested by a type III ANOVA and the random effect was tested by a likelihood ratio test comparing the model with and without the replicate lines. Both tests were performed using the library *lmerTest* package for R (Kuznetsova et al. 2017). If the selection effect was significant, *a posteriori* comparisons were performed using false discovery rate adjustment implemented in the *emmeans* package for R (Lenth et al. 2018).

Knockdown times were also used to plot the survival curves based on the Kaplan-Meier formula using the *survfit* function implemented in the *survival* package for R (Therneau 2023).

### Thermal death time curves (TDT)

Average knockdown times were calculated for each sex, replicate lines, and selection treatment combination (Table S1). These values were regressed against the assay temperatures according to Equation 1 (Rezende et al. 2014):

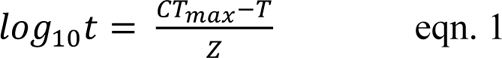

where *T* is the assay static temperature (°C), *CT*_max_ is the upper thermal limit (°C), *t* is the knockdown time (min), and *z* is the thermal sensitivity. These curves allowed the estimation of *CT*_max_ as the extrapolated temperature that would result in a knockdown time of log_10_ *t* = 0 (i.e., knockdown time at 1 min) and the estimation of the thermal sensitivity (*z* = –1/slope), where the lower *z* values, the higher the thermal sensitivity.

Using equation 1, 24 TDT curves (2 sexes × 3 replicate lines × 4 selection protocols) were fitted, from which *CT*_max_ and *z* values were estimated as described above. A linear model with ramping selection treatment (levels: fast-control, slow-control, fast-selection, and slow-selection lines), sex (levels: females and males), and their interaction was performed to evaluate their effects on *CT*_max_ and *z* values. TDT curve analysis did not include replicate lines as a random effect because only one *CT*_max_ and *z* value was estimated by each replicate line. Additionally, a linear mixed model with ramping selection (fixed effect with fast-control, slow-control, fast-selection, and slow-selection lines as levels), sex (fixed effect with females and males as levels), and replicate lines nested within the thermal selection (random effect with replicates 1, 2 and 3 as levels), and assay temperatures (as covariate) was fitted on the knockdown time using the *lmer* package for R.

### Desiccation and starvation resistance

To determine the lethal time at which 50% of flies of each vial were dead (LT_50_), a generalized linear model following a binomial distribution was fitted with the proportion of flies alive as the dependent variable and time as the predictor variable. The generalized linear model was run using the g*lm* function of the *lme4* package for R (Bates et al. 2015). The LT_50_ of each vial was then estimated using the function *dose.p* from the *MASS* package for R (Venables and Ripley 2002).

To estimate the median LT_50_ and the 95% confidence intervals for each selection treatment and sex, each LT_50_ was transformed into a survival object using the *Surv* and *survfit* functions of the *survival* package for R (Therneau 2023). This procedure also allowed to estimate the survival curves in each vial. Finally, to test the effect of selection treatment (levels: control, fast-selection, and slow-selection lines) and sex (levels: females and males) on desiccation and starvation resistance, a Cox proportional regression model was fitted with LT_50_ as the dependent variable, and selection protocol and sex as predictor variables. The Cox model was run using the *coxph* function of the *survival* package (Therneau 2023).

## RESULTS

### Knockdown temperature evolution

Knockdown temperature evolved in response to artificial selection for increased heat tolerance, regardless of the ramping assay protocol: the knockdown temperature was significantly higher in fast-ramping selected lines than in fast-ramping control lines (mean fast-ramping selected lines [95% CI] = 37.71 °C [37.63 – 37.78] and mean fast-ramping control lines [95% CI] = 37.23 °C [37.0 – 37.38]; *F*_1,4_ = 32.0, *P* = 0.005); and the knockdown temperature in slow-ramping selected lines was significantly higher than in slow-ramping control lines (mean slow-ramping selected lines [95% CI] = 35.48°C [35.41 – 35.55] and mean fast-ramping control lines [95% CI] = 34.97 °C [34.82 – 35.12]; *F*_1,4_ = 41.7, *P* = 0.003). These results were previously reported by Mesas et al. (2021) and are reported here to show that selected lines used in this study evolved higher thermal tolerance compared to control lines.

### Knockdown time evolution

As expected, the knockdown time decreased significantly as the assay temperatures increased (*F*_1,877_ = 649.1, *P* < 2×10^-16^). The mean knockdown time and 95% CI for each static assay are as follows: 35° C = 33.77 min [32.1 – 35.5]; 36° C = 16.98 min [16.1 – 17.9]; 37° C = 8.84 min [8.4 – 9.3]; and 38° C = 6.78 min [6.3 – 7.0].

Knockdown times differed significantly between selection treatments when flies were assayed at 36 and 37°C (Table 1; Table S2; Fig. 1). At these temperatures, slow and fast selected lines showed higher heat tolerance than slow and fast control lines (Table S1-S3; Fig. 1C, E). Also, fast-selected lines showed a higher heat tolerance than slow-selected lines in flies assayed at 37°C but not at 36°C (Table S1-S3; Fig. 1C, E), whereas fast and slow control lines did not differ (Table S3; Fig 1). On the other hand, replicate lines had no significant effect on knockdown time, indicating consistent evolutionary responses within each selection and control treatment (Table S2). Concerning sex, females showed a higher thermal tolerance than males but only when flies were assayed at 35 and 38°C (Table 1; Fig. 1B, H). Finally, non-significant interactions between selection and sex were found for all assay temperatures (Table 1).

**Figure 1.**
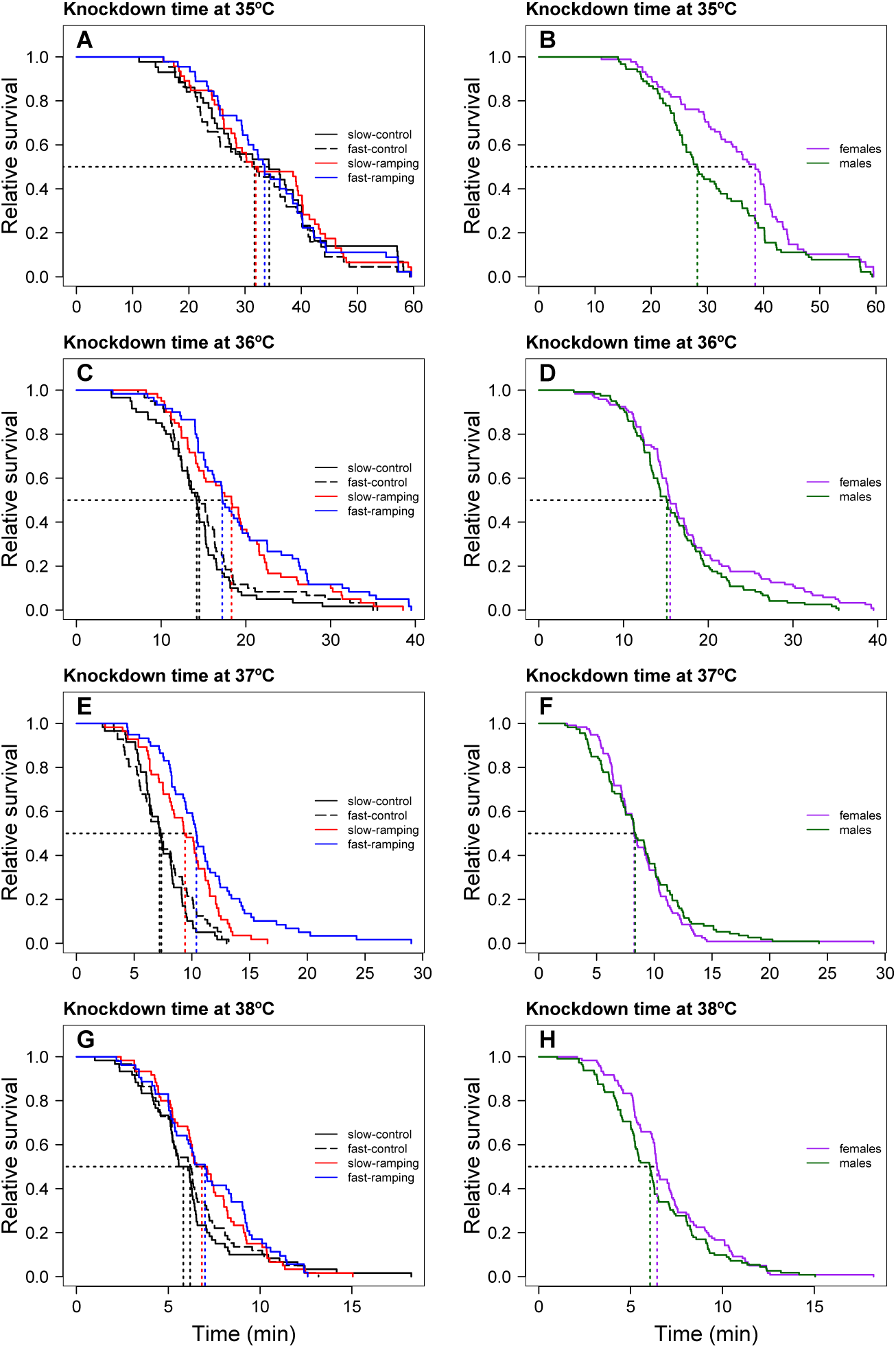
Heat-induced mortality in *Drosophila subobscura* flies assayed at four static temperatures. Left panels show the heat knockdown time of slow-ramping control (solid black line), fast-ramping control (dashed black line), slow-ramping selection (red line), and fast-ramping selection lines (blue lines). The right panels show the heat knockdown time of female (purple line) and male (green line) flies. Dotted lines indicate the median knockdown time for each selection protocol (left panels) and sex (right panels).

**Table 1.** Mixed linear effect model for the knockdown time of *Drosophila subobscura* assayed at four static temperature assays. For simplicity, results for the random effect (replicate lines) are not shown because they were not statistically significant (see Materials and Methods). Significant effects (P values < 0.05) are indicated in boldface type.

| Knockdown time | Selection | Sex | Selection × Sex |
| --- | --- | --- | --- |
| Static assay at 35°C | $F_{3,170} = 0.62$<br>$P = 0.60$ | $F_{1,170} = 8.64$<br>**$P = 0.004$** | $F_{3,170} = 0.64$<br>$P = 0.59$ |
| Static assay at 36°C | $F_{3,232} = 9.86$<br>**$P = 3.8 \times 10^{-6}$** | $F_{1,232} = 2.65$<br>$P = 0.10$ | $F_{3,232} = 0.74$<br>$P = 0.53$ |
| Static assay at 37°C | $F_{3,222} = 18.39$<br>**$P = 1.1 \times 10^{-10}$** | $F_{1,222} = 0.001$<br>$P = 0.97$ | $F_{3,222} = 2.05$<br>$P = 0.11$ |
| Static assay at 38°C | $F_{3,224} = 1.93$<br>$P = 0.13$ | $F_{1,224} = 4.63$<br>**$P = 0.032$** | $F_{3,224} = 2.44$<br>$P = 0.07$ |

### TDT curves evolution

Linear regressions between log_10_(LT_50_) and assay temperatures enabled the estimation of 24 TDT curves (4 selection treatments × 3 replicate lines × 2 sexes) with high coefficients of determination (mean *R*^2^ = 0.946, range: 0.820 – 0.989; Table S4), confirming that heat knockdown time is linearly related to stressful sublethal temperatures. From these TDT curves, the mean CT_max_ [95% CI] was 41.21°C [41.02 – 41.41], and the mean *z* [95% CI] was 4.18°C [4.03 – 4.32]. CT_max_ were significantly different between selection treatments (F_3,20_ = 4.46, *P* = 0.015; Fig. 2A). A post hoc analysis showed that fast-ramping selected and slow-ramping control lines were significantly different in their CT_max_ values (t_20_ = 3.195, *P* = 0.02). In contrast, fast and slow control lines had similar CT_max_ values (t_20_ = 0.911, *P* = 0.80). Thus, when control lines are pooled, CT_max_ still differs between selection treatments (F_2,18_ = 6.69, *P* = 0.007), with fast-ramping (mean CT_max_ [95% CI] = 41.55 °C [41.2 – 41.9]) and slow-ramping selected lines (mean CT_max_ [95% CI] = 41.43 °C [41.1 – 41.8]) had higher CT_max_ than control lines (mean CT_max_ [95% CI] = 40.94 °C [40.7 – 41.2]) (t_18_ = 3.27, *P* = 0.01 and t_18_ = 2.64, *P* = 0.04, respectively). CT_max_ was not different between the selected lines (t_18_ = 0.54, *P* = 0.85). On the other hand, sex and the interaction between selection treatments and sex had no significant effect on CT_max_ (F_1,18_ = 0.004, *P* = 0.95 and F_3,18_ = 2.11, *P* = 0.15, respectively). Regarding *z* (i.e., thermal sensitivity), it shows no significant effects of selection treatments (F_3,16_ = 0.91, *P* = 0.46; Fig. 2), sex (F_1,16_ = 1.30, *P* = 0.27), nor the interaction between selection treatments and sex (F_3,16_ = 2.23, *P* = 0.12). In summary, the evolution of a higher CT_max_ is not associated with an evolutionary change in thermal sensitivity (Fig. 2B). Indeed, the relationship between CT_max_ and *z* did not change with the evolution of increasing thermal tolerance (*r*_control-lines_ = 0.979 and *r*_selected-lines_ = 0.929; Z-test = 0.76, *P* = 0.45). Additionally, using a linear mixed model with the assay temperature as a covariate, this result was corroborated by a non-significant interaction between selection treatment and assay temperature (F_3,865_ = 0.30, *P* = 0.82).

**Figure 2.**
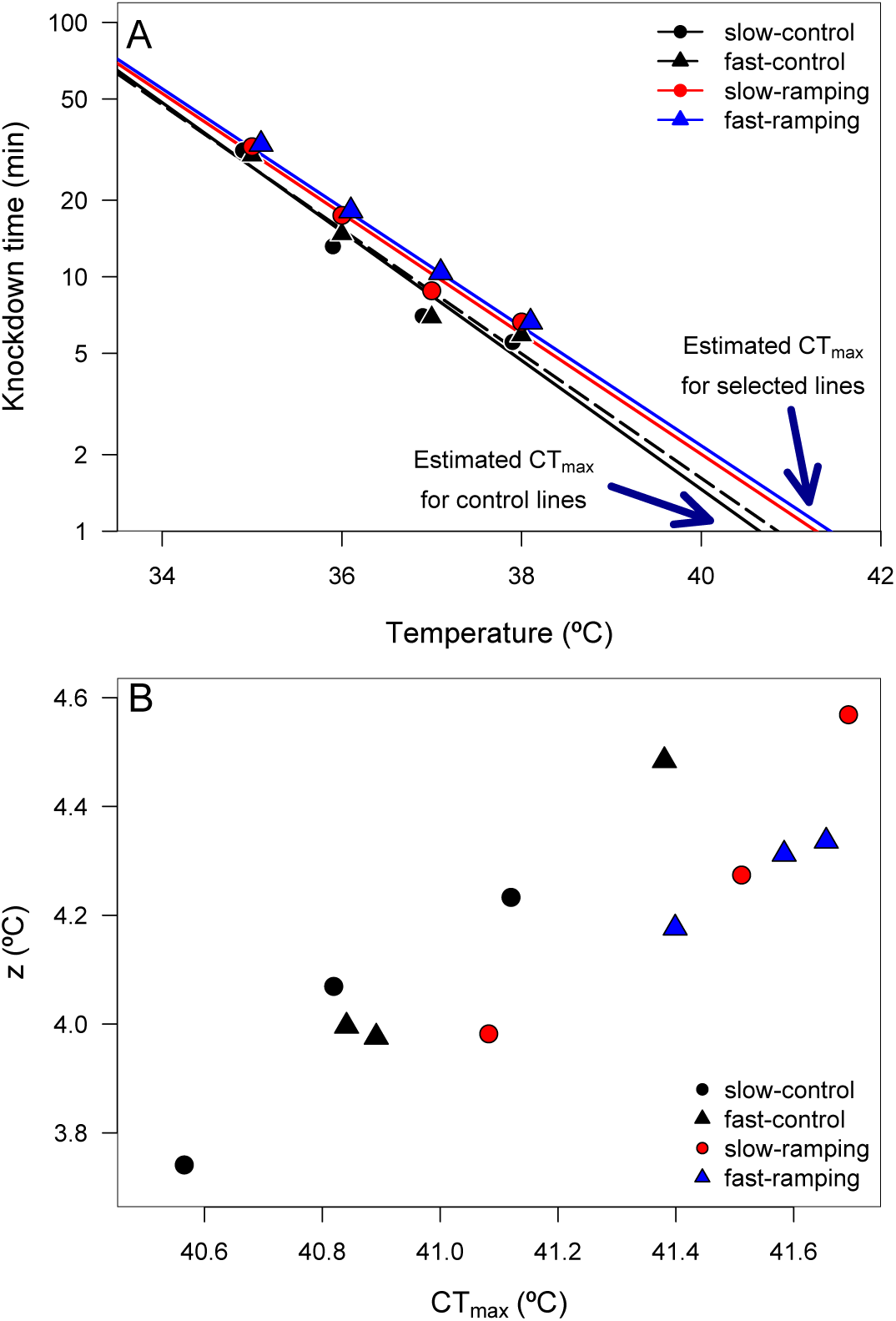
(A) Thermal death curves for control (black solid and dashed lines) and selected (red and blue lines) lines for increasing heat tolerance in *Drosophila subobscura*. Symbols represent the average knockdown time at the different assay temperatures. Each symbol represents the average knockdown time for each replicate line for each thermal regime: slow-control (black circle), fast-control (black triangle), slow-ramping (red circle), and fast-ramping (blue triangle). (B) Relationship between CT_max_ and z for slow-ramping control (solid black line), fast-ramping control (dashed black line), slow-ramping selection (red line), and fast-ramping selection lines (blue lines). Each symbol represents the CT_max_ and z estimated for each replicate line.

### Desiccation resistance evolution

Survival analysis showed a significant effect of sex and selection treatment on desiccation resistance, but not for the interaction of the two effects (Table S5). Males showed a higher risk of desiccation than female flies (hazard ratio = 7.11, *P* < 2×10^-7^; Fig. 3). Females showed a significant difference between selected and control lines (LTR: ×^2^_2_ = 6.72, *P* = 0.03; Fig. 3A). Specifically, females of the slow-ramping selection lines showed a higher desiccation resistance than females of the control lines (hazard ratio = 0.42, *P* = 0.009), whereas females of the fast-ramping selection and control lines showed similar desiccation risk (hazard ratio = 0.56, *P* = 0.072). On the other hand, males showed no differences in desiccation resistance between selected and control lines (LTR: *χ*^2^2 = 1.88, *P* = 0.4; Fig. 3B).

**Figure 3.**
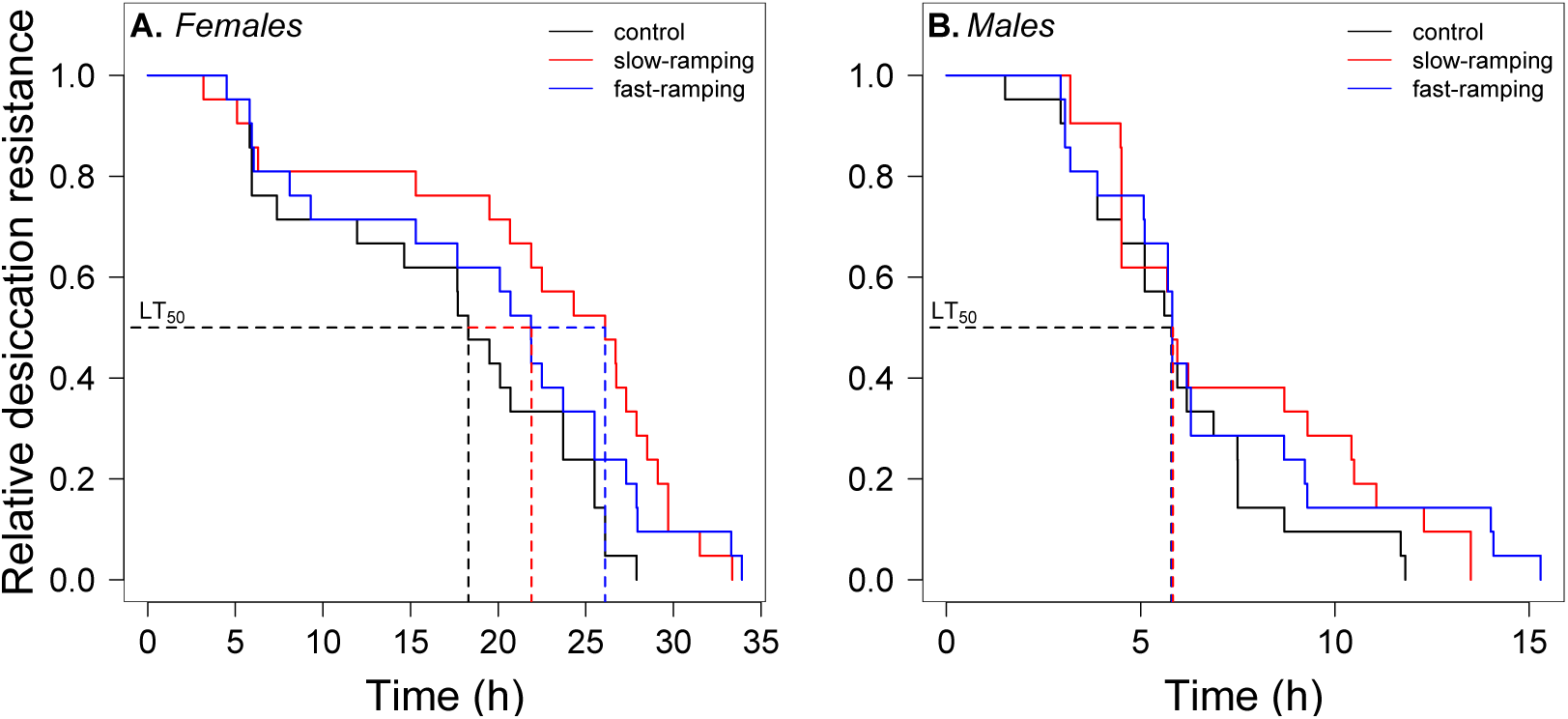
Desiccation survival curves of (A) females and (B) males from control (black line), slow-ramping selection (red line), and fast-ramping selection lines (blue lines) of *Drosophila subobscura*. Dashed lines indicate the median mortality time for each selection protocol (pooled replicate cages).

### Starvation resistance evolution

A significant effect of sex, selection treatment, and the interaction between the two effects on starvation resistance was found in the survival analysis (Table S6). Males had a higher risk of starvation than female flies (hazard ratio = 22.75, *P* < 1×10^-16^; Fig. 4). In female flies (Fig. 4A), fast-ramping selection and slow-ramping selection lines showed a higher starvation risk than control lines (hazard ratio = 2.37, *P* = 0.009; and hazard ratio = 2.20, *P* = 0.014, respectively). In contrast, male flies had an opposite pattern (Fig. 4B): slow-ramping selection lines had a lower starvation risk than control lines (hazard ratio = 0.50, *P* = 0.03), but nonsignificant differences were found between fast-ramping selection and control lines (hazard ratio = 0.64, *P* = 0.16).

**Figure 4.**
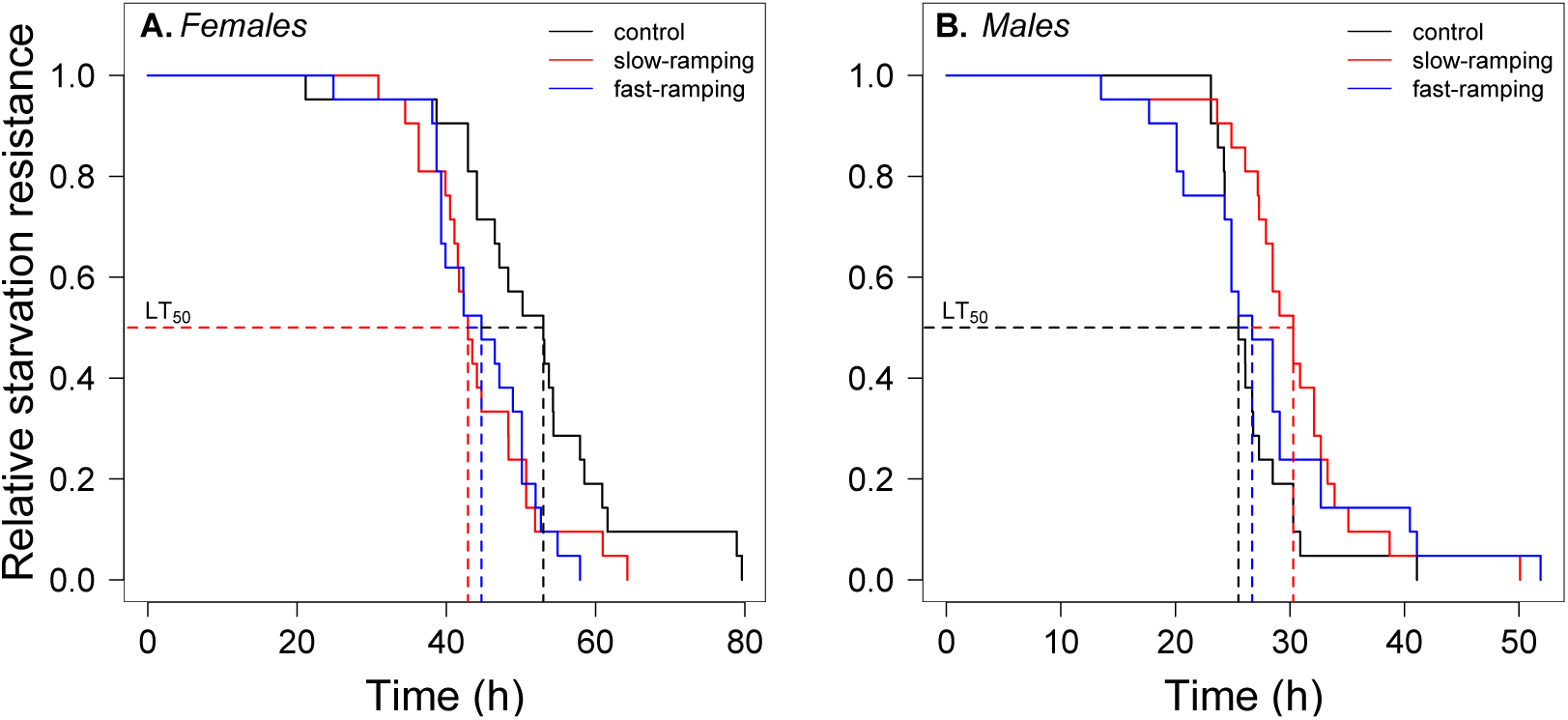
Starvation survival curves of (A) females and (B) males from control (black line), slow-ramping selection (red line), and fast-ramping selection lines (blue lines) of *Drosophila subobscura*. Dashed lines indicate the median mortality time for each selection protocol (pooled replicate cages).

## Discussion

Studying the evolutionary responses of thermal limits is key to understanding the adaptive responses and evolutionary constraints to global warming. Cross-tolerance studies can provide valuable information on the evolutionary response to multiple environmental stressors. Cross-tolerance evolution has been reported among different resistance traits (Hoffmann and Parsons 1993; Bubliy and Loeschcke 2005; Stazione et al. 2020; Singh et al. 2022), but the magnitude of the evolutionary response could be explained by the trait under direct selection or the stress intensity (Gerken et al. 2016). Here, artificial selection for heat tolerance (i.e., knockdown temperature) resulted in correlated responses in heat knockdown time, the thermal tolerance landscape (TDT curves), desiccation resistance, and starvation resistance. However, these responses depended on the intensity of thermal selection and sex, suggesting that the evolutionary response to tolerate higher temperatures also confers partial tolerance to other stresses such as desiccation and starvation.

Different approaches to measuring the upper thermal limit of ectotherms produce different genetic and phenotypic estimates. Fast-ramping assays generally estimate higher upper thermal limits and higher heritabilities than slow-ramping assays (Chown et al. 2009; Rezende et al. 2011). For instance, the heritability of thermal tolerance was 0.13 for fast assays and 0.08 for slow assays in *D. subobscura* (Castañeda et al. 2019). Because heritability is commonly used as a predictor of the evolutionary response of a trait to natural or artificial selection, the evolutionary response of heat tolerance would be expected to depend on the ramping rate used during selection.

However, previous work did not support this prediction for *D. subobscura*, finding that the evolution of heat tolerance was independent of the ramping rate (Mesas et al. 2021), but the correlated responses of the thermal performance curves or the energy metabolism depended on the intensity of the thermal selection (Mesas et al. 2021; Mesas and Castañeda 2023). In the present study, the evolution of knockdown temperature (e.g., heat tolerance measured in dynamic assays) induced a correlated response on the heat knockdown time (e.g., heat tolerance measured in static assays) when it was assayed at intermediate temperatures (36 and 37°C), but not at less or more extreme assay temperatures (35 and 38°C). These findings can be explained because stress tolerance at 35°C should depend on the physiological state of the organism during prolonged thermal assays (e.g., availability of energy resources; see Rezende et al. 2011, but also see Overgaard et al. 2012) and not only on heat tolerance, whereas heat tolerance at 38°C could be limited by physical properties of ectotherms (e.g., protein denaturation, membrane permeability). However, a previous study found a clinal pattern for heat tolerance in *D. subobscura* only for flies assayed in static assays (specifically at 38°C), but this clinal pattern was not detected using ramping assays (Castañeda et al. 2015). Differences between these two studies could be explained by the number of generations under thermal selection, which could result in a different evolutionary response of heat tolerance. According to Begon (1976), *D. subobscura* can have between 4 and 6 generations per year, which makes it possible to estimate about 125 generations of selection from the introduction of *D. subobcura* in Chile until the study by Castañeda et al. (2015). On the other hand, the type of selection is completely different between the two studies (e.g., natural versus artificial selection), which could lead to various evolutionary outcomes. In any case, beyond these results from specific thermal assays, these findings support the idea that (1) the use of a single static temperature would miss genetic or phenotypic effects on heat tolerance, and (2) unifying several knockdown time estimates into a single approach (TDT curves) should be necessary to elucidate genetic and phenotypic patterns of heat tolerance in ectotherms (Rezende et al. 2014; Jørgensen et al. 2021).

TDT curves evolved in response to heat tolerance selection in *D. subobscura*. TDT curves showed that fast- and slow-ramping selected lines evolved higher CT_max_ than control lines (ΔCT_max_ = 0.49 °C). This differential CT_max_ value is slightly lower than the population differences (0.9°C) observed between the lowest and highest latitude populations (∼8 latitudinal degrees) of *D. subobscura* studied by Castañeda et al. (2015) and even lower than the CT_max_ variation reported among *Drosophila* species (Jørgensen et al. 2019; Alruiz et al. 2022). On the other hand, although CT_max_ and z (i.e., thermal sensitivity) are phenotypically correlated (see Castañeda et al. 2015; Molina et al. 2023), the evolutionary increase in CT_max_ was not associated with a correlated response in thermal sensitivity (z). This result suggests that both thermal parameters are not genetically constrained, but further evidence from quantitative genetic studies is needed to assess the genetic association between CT_max_ and z. A caveat for this finding could be related to the fact that thermal selection for heat tolerance was carried out over 16 generations, followed by 7 generations of relaxed selection (i.e., no selection). However, previous evidence suggests that differences in heat tolerance between control and selected lines were consistent between generations 16 and 25 (Mesas et al. 2021). Indeed, Passananti et al. (2004) also reported that phenotypic values did not change after 35 generations of relaxed selection in desiccation-selected populations of *D. melanogaster*.

It was expected that flies selected for higher heat tolerance using slow-ramping rate protocols would exhibit greater desiccation and starvation resistance than flies selected using fast-ramping selection protocols. This is because flies assayed for heat tolerance in long assays are also exposed to desiccation and starvation stress (Santos et al. 2012). This study provides partial support for this hypothesis. First, slow-ramping selected lines evolved a higher desiccation resistance than control and fast-ramping selected lines. However, this was only observed in female flies, while males of the different selection treatments did not show any difference in desiccation resistance. On the other hand, starvation resistance evolved in opposite directions depending on sex: females of the fast-ramping and slow-ramping selected lines showed lower starvation resistance than females of the control lines, whereas males of the slow-ramping selected lines showed higher starvation resistance than males of the control and fast-ramping selected lines. Differential evolutionary responses between the sexes could be due to heat thermal selection only being applied to females, which could have exacerbated the evolutionary responses of female flies. However, previous studies that artificially selected exaggerated male traits also found fitness consequences in females (Harano et al. 2010). Differential evolutionary responses between females and males can then be explained by sexually antagonistic selection on genetically correlated traits (Eyer et al. 2019; Fanara et al. 2023). Kwan et al. (2008) reported that desiccation-selected females had higher desiccation resistance than desiccation-selected males (see also Chippindale et al. 2004), which can be explained by males using resources at a faster rate than females (e.g., males lose weight, water, and metabolites faster than females). Sexual dimorphism in stress resistance traits has been mainly explained by differences in cuticular composition, resource storage, and energy conservation between the sexes (Schwasinger-Schmidt et al. 2012; Rusuwa et al. 2022). Although energy content was not measured here, Mesas and Castañeda (2023) found that body mass and metabolic rate were similar between control and heat-tolerance selected lines of *D. subobscura*, suggesting that neither resource storage nor energy conservation explains the sex-dependent correlated response for stress resistance traits. However, the same study found that heat-tolerance selected lines had higher fecundity than control lines, whereas previous studies have found negative associations between fecundity and starvation resistance in *D. melanogaster* (Bubliy and Loeschcke 2005; Kalra et al. 2017). Then, the decrease in starvation resistance in females of the heat-selected lines could be related to increased fecundity, which is consistent with the reported trade-off between stress resistance traits and life-history traits (van Noordwijk and de Jong 1986; Rion and Kawecki 2007).

In conclusion, the present study shows that heat tolerance evolution is associated with evolutionary responses in other stress resistance traits, which could be explained by pleiotropic effects or linkage disequilibrium among the traits evaluated. However, further evidence (e.g., quantitative genetic or genome-wide analysis studies) is needed to elucidate the genetic basis of the cross-tolerance evolution in *D. subobscura*. In addition, this study provides evidence for rapid evolutionary responses in ectotherms mediated by thermal selection, but the evolutionary outcomes depend on the intensity of the thermal stress (Mesas and Castañeda 2023) and sex (Rogell et al. 2014; Rusuwa et al. 2022). This study also highlights the importance of *D. subobscura* as a suitable model to study thermal adaptation mediated by natural selection (Huey 2000; Gilchrist et al. 2008; Castañeda et al. 2013, 2015), and laboratory selection (Santos et al. 2005, Santos et al. 2021; Simões et al. 2017; Mesas et al. 2021; Mesas and Castañeda 2023). In addition, this study highlights the relevance of experimental evolutionary studies for understanding the adaptive responses to climate change (Mitchell and Whitney 2018; Brennan et al. 2022; Kelly 2022). Finally, these results suggest that ectotherms may evolve in response to climate warming, but evolutionary responses may differ between sexes and/or the warming rates experienced by natural populations, which may make it difficult to propose general trends in the fate of ectotherms in a changing world where temperature is not the only driver of climate change, but species are also expected to be exposed to changes in precipitation patterns and food availability.

## Acknowledgments

I thank Andres Mesas, Angélica Jaramillo, Julio Figueroa, and Jaiber Solano for their help with fly maintenance, experiments, and data management. I also thank Enrico Rezende and Mauro Santos for their valuable suggestions and advice while designing the artificial selection experiment. A preprint version of this article has been peer-reviewed and recommended by PCIEvolBiol (https://doi.org/10.24072/pci.evolbiol.100709).

## Funding

This work was funded by the Fondo Nacional de Desarrollo Científico y Tecnológico (FONDECYT) [grant number 1140066] – Chile. Currently, L.E. Castañeda is supported by the Agencia Nacional de Invertigación y Desarrollo (ANID) through the grants Anillo ATE230025, FOVI220194, and FOVI 230149.

## Data availability

Data and scripts are available at https://doi.org/10.6084/m9.figshare.24085107.v5

## Conflict of interest disclosure

The author declares having no financial conflicts in relation to the content of the article.

## Notes

https://doi.org/10.6084/m9.figshare.24085107.v5

### Competing Interest Statement

The authors have declared no competing interest.

### Summary of Updates

The final version including information of the review process.

