## Supplementary material for "Cross-tolerance evolution is driven by selection on heat tolerance in *Drosophila subobscura*"

**Figure S1**. Diagram of the experimental design.

**Table S1**. Descriptive statistics of knockdown time of *Drosophila subobscura*.

**Table S2**. Results of the mixed-linear models on the knockdown time of *Drosophila subobscura.*

**Table S3**. Tukey's contrast analysis for the knockdown time of *Drosophila subobscura*.

**Table S4**. Thermal-death-time (TDT) curves for *Drosophila subobscura*.

**Table S5**. Results of the desiccation survival of *Drosophila subobscura*.

**Table S6**. Results of the starvation survival of *Drosophila subobscura*.

**Figure S1.** Diagram of the experimental design: 100 isofemale lines from *Drosophila subobscura* were used to establish an outbred population. The F1 of these isofemale lines were transferred to a population cage and the F2 flies were divided into three replicates: R1, R2, and R3. After 3 generations, each population cage was divided into four population cages, which were assigned to four different artificial selection protocols in triplicate: fast-ramping selection, fast-ramping control, slow-ramping selection, and slow-ramping control lines. During 16 generations, heat tolerance was selected for 33% highest values of knockdown temperature using two different selection protocols: slow ramping rate (0.08 ºC/min) and fast ramping rate (0.4 ºC/min). Traits were evaluated at generation 23 and 24.

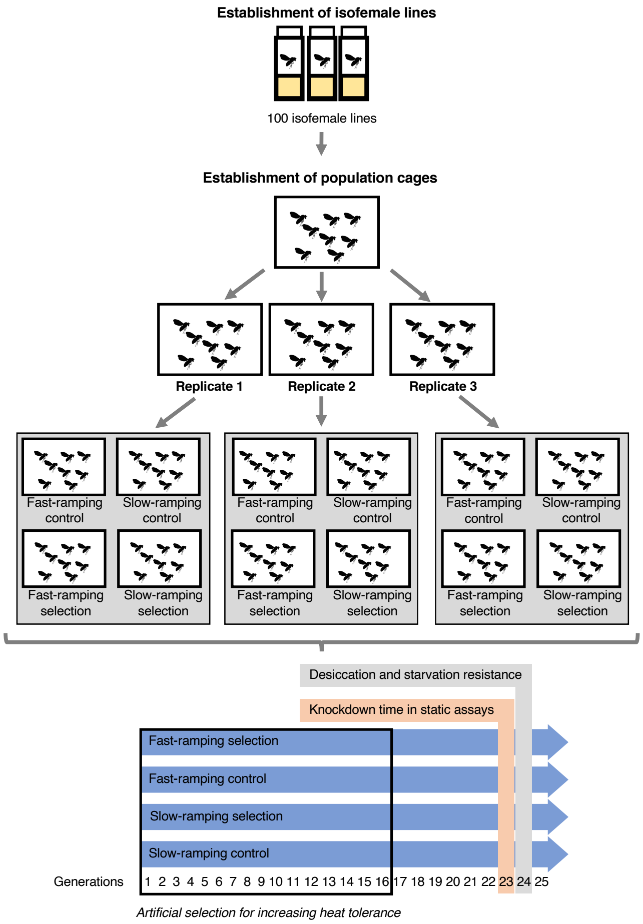

**Table S1**. Mean (SD) knockdown time of *Drosophila subobscura* flies assayed at four static temperatures. Values are organized by sex, selection protocol, and replicate cage.

| Selection protocol | Replicated cage | Females | | | | Males | | | |
|---|---|---|---|---|---|---|---|---|---|
| | | Knockdown time at 35ºC (min) | Knockdown time at 36ºC (min) | Knockdown time at 37ºC (min) | Knockdown time at 38ºC (min) | Knockdown time at 35ºC (min) | Knockdown time at 36ºC (min) | Knockdown time at 37ºC (min) | Knockdown time at 38ºC (min) |
| Fast-ramping control | R1 | 28.91 (1.48) | 14.66 (1.20) | 7.34 (1.31) | 6.17 (1.34) | 25.39 (1.56) | 14.09 (1.42) | 7.95 (1.36) | 5.73 (1.77) |
| | R2 | 33.71 (1.26) | 13.86 (1.49) | 6.94 (1.39) | 5.95 (1.44) | 30.02 (1.25) | 15.00 (1.41) | 6.49 (1.58) | 5.55 (1.51) |
| | R3 | 30.13 (1.54) | 15.51 (1.36) | 7.88 (1.49) | 6.75 (1.35) | 33.64 (1.48) | 15.38 (1.46) | 5.60 (1.55) | 5.33 (1.53) |
| Slow-ramping control | R1 | 34.30 (1.49) | 15.80 (1.43) | 6.91 (1.25) | 5.74 (1.26) | 25.43 (1.42) | 12.07 (1.34) | 5.86 (1.58) | 5.65 (1.50) |
| | R2 | 38.16 (1.51) | 10.86 (1.46) | 7.31 (1.18) | 5.82 (1.80) | 28.15 (1.60) | 13.75 (1.26) | 7.25 (1.34) | 5.63 (2.14) |
| | R3 | 33.84 (1.67) | 13.80 (1.64) | 7.09 (1.59) | 5.42 (1.44) | 31.18 (1.37) | 13.34 (1.63) | 7.62 (1.34) | 5.04 (1.48) |
| Fast-ramping selection | R1 | 31.83 (1.35) | 22.08 (1.55) | 10.52 (1.64) | 7.64 (1.41) | 30.56 (1.58) | 16.81 (1.43) | 11.12 (1.52) | 5.58 (1.24) |
| | R2 | 37.69 (1.21) | 18.30 (1.43) | 9.02 (1.30) | 7.87 (1.34) | 39.11 (1.37) | 19.14 (1.30) | 11.94 (1.33) | 4.91 (1.69) |
| | R3 | 37.45 (1.46) | 17.20 (1.89) | 9.17 (1.38) | 7.81 (1.57) | 32.45 (1.21) | 16.11 (1.39) | 11.08 (1.54) | 5.69 (1.69) |
| Slow-ramping selection | R1 | 34.72 (1.29) | 19.95 (1.42) | 9.48 (1.24) | 6.54 (1.44) | 29.76 (1.50) | 18.17 (1.55) | 9.73 (1.53) | 7.11 (1.22) |
| | R2 | 32.06 (1.50) | 18.09 (1.36) | 10.01 (1.30) | 7.09 (1.37) | 26.38 (1.44) | 16.98 (1.34) | 6.49 (1.59) | 6.82 (1.55) |
| | R3 | 41.97 (1.29) | 18.40 (1.52) | 8.36 (1.44) | 6.53 (1.51) | 33.08 (1.34) | 13.81 (1.33) | 9.34 (1.42) | 5.85 (1.63) |

**Table S2**. Results of the mixed-linear models on the knockdown time of *Drosophila subobscura*. Fixed effects were tested by a type III ANOVA, and the random effect was tested by a likelihood ratio test comparing the model with and without the replicate lines. Response variables were squared root transformed. Significant effects ($P < 0.05$) are indicated in bold.

**Knockdown time at 35ºC**

| *Fixed effect* | *SS* | $DF_{num}, DF_{den}$ | *F* | *P value* |
|---|---|---|---|---|
| Selection | 1.82 | 3,170 | 0.62 | 0.602 |
| Sex | 8.44 | 1,170 | 8.64 | **0.004** |
| Selection × Sex | 1.89 | 3,170 | 0.64 | 0.588 |
| *Random effect* | *Variance* | *Likelihood ratio test (df=1)* | | *P value* |
| Replicate(Selection) | 0.0000 | 0 | | 1 |
| Error | 0.9767 | | | |

**Knockdown time at 36ºC**

| *Fixed effect* | *SS* | $DF_{num}, DF_{den}$ | *F* | *P value* |
|---|---|---|---|---|
| Selection | 16.68 | 3,232 | 9.86 | $\mathbf{3.8 \times 10^{-6}}$ |
| Sex | 1.49 | 1,232 | 2.65 | 0.10 |
| Selection × Sex | 1.25 | 3,232 | 0.74 | 0.53 |
| *Random effect* | *Variance* | *Likelihood ratio test (df=1)* | | *P value* |
| Replicate(Selection) | 0.0000 | 0 | | 1 |
| Error | 0.5639 | | | |

**Knockdown time at 37ºC**

| *Fixed effect* | *SS* | $DF_{num}, DF_{den}$ | *F* | *P value* |
|---|---|---|---|---|
| Selection | 14.96 | 3,223 | 18.75 | $\mathbf{7.0 \times 10^{-11}}$ |
| Sex | 0.002 | 1,223 | 0.009 | 0.93 |
| Selection × Sex | 1.58 | 3,223 | 1.99 | 0.12 |
| *Random effect* | *Variance* | *Likelihood ratio test (df=1)* | | *P value* |
| Replicate(Selection) | 0.0000 | 0 | | 1 |
| Error | 0.2656 | | | |

**Knockdown time at 38ºC**

| *Fixed effect* | *SS* | $DF_{num}, DF_{den}$ | *F* | *P value* |
|---|---|---|---|---|
| Selection | 1.69 | 3,224 | 2.27 | 0.08 |
| Sex | 1.42 | 1,224 | 5.70 | **0.02** |
| Selection × Sex | 1.74 | 3,224 | 2.33 | 0.08 |
| *Random effect* | *Variance* | *Likelihood ratio test (df=1)* | | *P value* |
| Replicate(Selection) | 0.0000 | 0 | | 1 |
| Error | 0.2485 | | | |

**Table S3.** Tukey's contrast analysis for the knockdown time of *Drosophila subobscura* assayed in four static temperature assays. P values were corrected using the false discovery rate method. Significant differences ($P < 0.05$) are indicated in bold.

**Knockdown time at 35ºC**

No selection effect

**Knockdown time at 36ºC**

| *Contrasts* | *df* | *t* | *P value* |
|---|---|---|---|
| slow-control vs. fast-control | 8 | -1.426 | 0.230 |
| slow-control vs. slow-selected | 8 | -3.994 | **0.012** |
| slow-control vs. fast-selected | 8 | -4.773 | **0.008** |
| fast-control vs. slow-selected | 8 | -2.568 | 0.050 |
| fast-control vs. fast-selected | 8 | -3.347 | **0.020** |
| slow-selected vs. fast-selected | 8 | -0.779 | 0.458 |

**Knockdown time at 37ºC**

| *Contrasts* | *df* | *t* | *P value* |
|---|---|---|---|
| slow-control vs. fast-control | 7.91 | -0.010 | 0.992 |
| slow-control vs. slow-selected | 8.06 | -3.432 | **0.013** |
| slow-control vs. fast-selected | 7.64 | -6.368 | **0.001** |
| fast-control vs. slow-selected | 8.33 | -3.392 | **0.013** |
| fast-control vs. fast-selected | 7.91 | -6.302 | **0.001** |
| slow-selected vs. fast-selected | 8.06 | -2.841 | **0.026** |

**Knockdown time at 38ºC**

No selection effect

**Table S4**. Thermal-death-time (TDT) parameters calculated from heat tolerance measurements for *Drosophila subobscura*.

| Selection regimen | Replicate | Sex | TDT curve | $CT_{max}$ (ºC) | *z* (ºC) | $r^2$ | $Q_{10}$ |
|---|---|---|---|---|---|---|---|
| Fast-ramping control | R1 | females | $\log_{10} t = 9.6218 – 0.2338\ T$ | 41.15 | 4.28 | 0.9425 | 217.84 |
| | R2 | females | $\log_{10} t = 10.3173 – 0.2527\ T$ | 40.83 | 3.96 | 0.9297 | 336.39 |
| | R3 | females | $\log_{10} t = 9.4471 – 0.2281\ T$ | 41.42 | 4.38 | 0.9469 | 190.81 |
| | R1 | males | $\log_{10} t = 8.8691 – 0.2132\ T$ | 41.61 | 4.69 | 0.9637 | 135.36 |
| | R2 | males | $\log_{10} t = 10.1218 – 0.2478\ T$ | 40.85 | 4.03 | 0.9484 | 300.91 |
| | R3 | males | $\log_{10} t = 11.3142 – 0.2803\ T$ | 40.36 | 3.57 | 0.9147 | 635.93 |
| Slow-ramping control | R1 | females | $\log_{10} t = 11.1809 – 0.2761\ T$ | 40.49 | 3.62 | 0.9455 | 577.29 |
| | R2 | females | $\log_{10} t = 10.2409 – 0.2506\ T$ | 40.87 | 3.99 | 0.8199 | 320.36 |
| | R3 | females | $\log_{10} t = 11.0648 – 0.2731\ T$ | 40.52 | 3.66 | 0.9572 | 538.05 |
| | R1 | males | $\log_{10} t = 9.1095 – 0.2214\ T$ | 41.15 | 4.52 | 0.8985 | 163.61 |
| | R2 | males | $\log_{10} t = 9.2449 – 0.2235\ T$ | 41.37 | 4.47 | 0.9132 | 171.69 |
| | R3 | males | $\log_{10} t = 10.6311 – 0.2618\ T$ | 40.61 | 3.82 | 0.9760 | 414.62 |
| Fast-ramping selection | R1 | females | $\log_{10} t = 9.0865 – 0.2155\ T$ | 42.16 | 4.64 | 0.9789 | 143.04 |
| | R2 | females | $\log_{10} t = 9.6911 – 0.2329\ T$ | 41.60 | 4.29 | 0.9357 | 213.53 |
| | R3 | females | $\log_{10} t = 9.7291 – 0.2336\ T$ | 41.65 | 4.28 | 0.9337 | 216.83 |
| | R1 | males | $\log_{10} t = 10.2040 – 0.2479\ T$ | 41.15 | 4.03 | 0.9875 | 301.61 |
| | R2 | males | $\log_{10} t = 10.1460 – 0.2463\ T$ | 41.20 | 4.06 | 0.9783 | 290.34 |
| | R3 | males | $\log_{10} t = 9.5585 – 0.2302\ T$ | 41.52 | 4.34 | 0.9863 | 200.49 |

| | | | | | | | |
|---|---|---|---|---|---|---|---|
| Slow-ramping selection | R1 | females | $\log_{10} t = 10.2076 - 0.2474\ T$ | 41.25 | 4.04 | 0.9797 | 298.12 |
| | R2 | females | $\log_{10} t = 9.4324 - 0.2263\ T$ | 41.69 | 4.42 | 0.9840 | 183.12 |
| | R3 | females | $\log_{10} t = 11.0759 - 0.2711\ T$ | 40.86 | 3.69 | 0.9551 | 513.69 |
| | R1 | males | $\log_{10} t = 9.2689 - 0.2219\ T$ | 41.77 | 4.51 | 0.9895 | 165.59 |
| | R2 | males | $\log_{10} t = 9.8391 - 0.2120\ T$ | 41.70 | 4.72 | 0.8784 | 131.73 |
| | R3 | males | $\log_{10} t = 9.6621 - 0.2339\ T$ | 41.31 | 4.27 | 0.9567 | 218.37 |

**Table S5**. Results of the desiccation survival analysis testing the effect of selection protocol, sex, and their interaction in *Drosophila subobscura*. Significant effects ($P < 0.05$) are indicated in bold.

| **Effect** | **exp(coefficient)** | **z** | ***P value*** |
|---|---|---|---|
| Slow-ramping selection | 0.422 | –2.623 | **0.009** |
| Fast-ramping selection | 0.556 | –1.797 | 0.072 |
| Males | 7.108 | 5.242 | **$1.6 \times 10^{-7}$** |
| Slow-ramping selection – males | 1.773 | 1.264 | 0.206 |
| Fast-ramping selection –males | 1.391 | 0.731 | 0.465 |

| **Selection treatment** | **vials** | **Median (h)** | **95% CI (h)** |
|---|---|---|---|
| *Females* | | | |
| Control | 21 | 18.3 | 11.9 – 25.5 |
| Slow-ramping selection | 21 | 26.1 | 20.7 – 29.1 |
| Fast-ramping selection | 21 | 21.9 | 15.3 – 27.3 |
| *Males* | | | |
| Control | 21 | 5.81 | 4.51 – 7.50 |
| Slow-ramping selection | 21 | 5.81 | 4.51 – 10.49 |
| Fast-ramping selection | 21 | 5.81 | 5.10 – 9.23 |

| **Sex** | **vials** | **Median (h)** | **95% CI (h)** |
|---|---|---|---|
| Females | 63 | 21.87 | 19.5 – 25.5 |
| Males | 63 | 5.81 | 5.61 – 6.3 |

**Table S6**. Results of the starvation survival analysis testing the effect of selection protocol, sex, and their interaction in *Drosophila subobscura*. Significant effects ($P < 0.05$) are indicated in bold.

| Effect | exp(coefficient) | z | *P value* |
|---|---|---|---|
| Slow-ramping selection | 2.062 | 2.240 | **0.025** |
| Fast-ramping selection | 2.268 | 2.474 | **0.013** |
| Males | 22.749 | 8.212 | **$<2\times10^{-16}$** |
| Slow-ramping selection – males | 0.223 | –3.305 | **0.0009** |
| Fast-ramping selection –males | 0.218 | –3.273 | **0.001** |

| Selection treatment | vials | Median (h) | 95% CI (h) |
|---|---|---|---|
| *Females* | | | |
| Control | 21 | 53.0 | 46.5 – 58.5 |
| Slow-ramping selection | 21 | 42.9 | 41.1 – 50.7 |
| Fast-ramping selection | 21 | 44.7 | 39.3 – 50.1 |
| *Males* | | | |
| Control | 21 | 25.5 | 24.9 – 28.5 |
| Slow-ramping selection | 21 | 30.3 | 27.9 – 33.3 |
| Fast-ramping selection | 21 | 26.7 | 24.9 – 32.7 |

| Sex | vials | Median (h) | 95% CI (h) |
|---|---|---|---|
| Females | 63 | 46.5 | 42.9 – 50.1 |
| Males | 63 | 27.3 | 26.1 – 29.1 |
